# C57BL/6 substrain differences in inflammatory and neuropathic nociception and genetic mapping of a major quantitative trait locus underlying acute thermal nociception

**DOI:** 10.1101/413856

**Authors:** Camron D. Bryant, Deniz Bagdas, Lisa R. Goldberg, M. Tala Khalefa, M.Eric R. Reed, Stacey L. Kirkpatrick, Julia C. Kelliher, Melanie M. Chen, William E. Johnson, Megan K. Mulligan, M. Imad Damaj

## Abstract

Sensitivity to different pain modalities has a genetic basis that remains largely unknown. Employing closely related inbred mouse substrains can facilitate gene mapping of nociceptive behaviors in preclinical pain models. We previously reported enhanced sensitivity to acute thermal nociception in C57BL/6J (**B6J**) versus C57BL/6N (**B6N**) substrains. Here, we expanded on nociceptive phenotypes and observed an increase in formalin-induced inflammatory nociceptive behaviors and paw diameter in B6J versus B6N mice (Charles River Laboratories). No strain differences were observed in mechanical or thermal hypersensitivity or in edema following the Complete Freund’s Adjuvant (CFA) model of inflammatory pain, indicating specificity in the inflammatory nociceptive stimulus. In the chronic nerve constriction injury (CCI), a model of neuropathic pain, no strain differences were observed in baseline mechanical threshold or in mechanical hypersensitivity up to one month post-CCI. We replicated the enhanced thermal nociception in the 52.5°C hot plate test in B6J versus B6N mice from The Jackson Laboratory. Using a B6J x B6N-F2 cross (N=164), we mapped a major quantitative trait locus (**QTL**) underlying hot plate sensitivity to chromosome 7 that peaked at 26 Mb (LOD=3.81, p<0.01; 8.74 Mb-36.50 Mb) that was more pronounced in males. Genes containing expression QTLs (eQTLs) associated with the peak nociceptive marker that are implicated in pain and inflammation include *Ryr1, Cyp2a5, Pou2f2, Clip3, Sirt2, Actn4*, and *Ltbp4* (FDR < 0.05). Future studies involving positional cloning and gene editing will determine the quantitative trait gene(s) and potential pleiotropy of this locus across pain modalities.

## INTRODUCTION

Pain is defined by the International Association for the Study of Pain as an unpleasant sensory and or emotional experience associated with actual or potential tissue damage (http://www.iasp-pain.org) and is the number one reason patients contact their physician. Nociceptive pain is adaptive and signals potential or actual tissue damage and thus promotes avoidance and protective behavior. However, chronic pain (e.g., neuropathic pain) is maladaptive and significantly decrease quality of life. An estimated one-third of the global population suffers from chronic pain, including over 100 million people in the United States^(1)^. Both environmental and genetic factors contribute to the wide range of variability in pain sensitivity^(2, 3)^. Pain has multiple modalities, including thermal, inflammatory, mechanical hypersensitivity, and neuropathic pain. Each of these modalities are hypothesized to have largely separable genetic components^(4-6)^. Furthermore, pain has both peripheral and central nervous system components with both shared and divergent molecular and physiological functions. Estimates from preclinical models indicate that 30-80% of the variance in pain responses can be explained by genetic factors^(7)^ whereas heritability estimates from human experimental pain models range from 20 to 60%^(2, 7)^.

Human genome-wide association studies (GWAS) have had limited success in identifying replicable “pain genes”, due in large part to a lack of statistical power and a lack of a consistent definition of pain cases across cohorts^(1, 8)^. Quantitative trait locus (**QTL**) mapping is an unbiased, discovery-based (i.e, hypothesis-generating) approach to identifying polymorphic regions of the genome containing genetic variants underlying complex traits. QTL mapping in preclinical mammalian models of pain provides multiple advantages, including a greater ability to achieve the necessary statistical power and the ability to combine gene expression analysis of relevant tissue (e.g., PNS and CNS tissue) with behavioral QTL analysis to facilitate the identification of quantitative trait genes and functional variants^(9)^. QTL studies of nociceptive behaviors associated with pain models in mice can facilitate translation to humans^(1, 2)^. The *CACNG2* gene (coding for the gamma 2 subunit of a calcium channel) was first mapped in mice for nerve injury-induced autonomy, using scratching, biting, and licking as the heritable behavioral pain measures for mapping^(10)^. Haplotypes within *CACNG2* were subsequently associated with increased post-mastectomy neuropathic pain^(1)^. QTL mapping in mice also identified a non-synonymous SNP in *P2X7* (encoding the adenosine triphosphate–gated P2X7 purinergic receptor) that was associated with reduced allodynia in a spared nerve injury model^(11)^. *P2X7* was subsequently associated with post-mastectomy pain in women^(1)^. QTL mapping in mice also identified calcitonin gene-related peptide (*CGRP*) in thermal nociception^(12)^ and the *MC1R* (the gene encoding the melanocortin 1 receptor) in female-specific kappa opioid-induced antinociception – a finding that was subsequently confirmed in women^(13)^. As a final example, a QTL for inflammatory pain was mapped to *AVPR1A* (gene encoding the vasopressin 1A receptor) and in humans, a SNP in *AVPR1A* was subsequently associated with inflammatory pain and vasopressin receptor-mediated analgesia in non-stressed but not stressed men^(14)^.

Despite successes in translating pain genetics from mice to humans, gene mapping in mice remains a challenge for two main reasons. First, genetic complexity underlying a QTL can hinder gene identification. In comparison to the reference C57BL/6J (**B6J**) inbred strain, a majority of classical inbred strains contain five million or so variants (SNPs plus indels); in stark contrast, the closely related C57BL/6NJ (**B6NJ**) inbred substrain contains approximately 30,000 variants^(15, 16)^. Thus, the magnitude of genetic complexity underlying a QTL is decreased by orders of magnitude which facilitates the identification of causal genes and variants in these so called Reduced Complexity Crosses (**RCCs**)^(17-20)^. The second hurdle to gene identification concerns the low level of resolution of QTLs, especially in F2 crosses, where the confidence interval typically spans one-half of a chromosome and contains hundreds of genes and thousands of variants. While the use of RCCs does not provide immediate improvement of QTL resolution like other contemporary mouse populations and panels such as highly recombinant outbred stocks^(9)^, RCCs can facilitate the subsequent step of fine mapping due to the much simpler genetic architecture of the quantitative traits (typically monogenic inheritance on a nearly isogenic background) by permitting immediate backcrossing and phenotyping at each generation as new recombination events accumulate, thus, rapidly narrowing the interval^(20)^. We used C57BL/6 (B6) substrains to map the genetic basis of parental strain variation in binge eating between C57BL/6 substrains^(19)^. We identified a single QTL on mid-chromosome 11 that mapped to the same region identified for cocaine neurobehavioral sensitivity using the same RCC^(18)^. The region contains a missense mutation in the gene *Cyfip2* that could act as a gain-of-function allele. Accordingly, mice heterozygous for a null mutation in *Cyfip2* showed a normalization in binge eating toward a wild-type level, thus providing support for *Cyfip2* as a causal genetic factor underlying binge eating in B6 substrains^(18)^.

With respect to pain models, we previously identified behavioral differences in B6 substrains in two assays of acute thermal nociception, including the hot plate and tail withdrawal assays. In both cases, the B6J substrain showed enhanced nociceptive sensitivity relative to the B6N substrains as indicated by a reduction in latency to respond to the thermal nociceptive stimuli^(21)^. Subsequent studies replicated the increase in acute thermal nociceptive sensitivity in B6J relative to other B6N^(22)^. However, it is unknown whether enhanced nociceptive sensitivity in the B6J substrain extends to other nociceptive modalities besides acute thermal nociception. A recent report indicated no differences in mechanical sensitivity^(23)^. Furthermore, because genetic factors are hypothesized to underlie B6 substrain differences in nociceptive behaviors, in the second part of this study, we sought to map the genetic basis of acute thermal nociception in the hot plate assay^(21)^. To prioritize functional candidate genes for future gene editing, we used a historical B6J x B6NJ-F2 genomic dataset to report genes within the behavioral QTL for hot plate sensitivity that also possess a *cis*-expression QTL (eQTL) within striatal brain tissue, including a top differentially expressed candidate gene, the ryanodine receptor 1 (*Ryr1*). Genes within the locus whose expression is associated with both genetic variation and behavior are prioritized as candidate quantitative trait genes underlying thermal nociception.

## MATERIALS AND METHODS

### Mice

Adult (8–10 weeks of age at the beginning of experiments) female and male C57BL/6J (**B6J**) (The Jackson Laboratory; JAX - Bar Harbor, ME USA) and C57BL/6N mice (**B6N**) (Charles River Laboratories - Wilmington, MA USA) were used for phenotyping in the inflammatory and neuropathic pain models. Mice were housed in a 21°C humidity-controlled Association for Assessment and Accreditation of Laboratory Animal Care–approved animal care facility in Virginia Commonwealth University. Mice were housed in groups of four and had free access to food and water. The rooms were on a 12-hour light/dark cycle (lights on at 7:00 AM). All experiments were performed during the light cycle, and the study was approved by the Institutional Animal Care and Use Committee of Virginia Commonwealth University. All studies were carried out in accordance with the National Institutes of Health’s Guide for the Care and Use of Laboratory Animals.

For experiments involving hot plate testing and QTL analysis (Boston University School of Medicine; BUSM), C57BL/6J (**B6J**) and C57BL/NJ (**B6NJ**) mice were purchased from JAX at 7 weeks of age and were habituated in the vivarium one week prior to experimental testing that occurred next door. All behavioral testing was performed during the light phase of the 12 h light/dark cycle (0630 h/1830 h). Testing occurred between 0800 h and 1300 h. For QTL mapping, B6J females were crossed to B6NJ males to generate B6J x B6NJ-F1 offspring and B6J x B6NJ F1 mice were intercrossed to generate B6J x B6NJ F2 mice. Mice were 50-100 days old at the time of testing. All experiments were approved by the Institutional Animal Care and Use Committee of BUSM and completed in accordance with the National Institutes of Health’s Guide for the Care and Use of Laboratory Animals.

### Drugs

Complete Freund Adjuvant (CFA) was purchased from Sigma-Aldrich (St. Louis, MO). CFA was diluted with mineral oil and administered at a concentration of 10%. Formalin was purchased from Fisher Scientific (Waltham, MA) and diluted with distilled water to a 2.5% concentration. Formalin was prepared daily.

### Formalin test of inflammatory nociception

Female and male B6J and B6N mice (n=6/sex/group) were used to study formalin nociceptive behaviors. The formalin test was carried out in an open Plexiglas cage (29 × 19 × 13 cm each). Mice were allowed to acclimate for 5 min in the test cage prior to injection. Each animal was injected with 20 μL of formalin (2.5%) to the dorsal surface of the right hind paw. Mice were observed from 0 to 5 min (early phase) and 20 to 45 min (late phase) post-formalin injection. The amount of time spent licking the injected paw was recorded with a digital stopwatch. Paw diameter (see measurement of paw edema) was also determined before and 1 h after formalin injection.

### Complete Freund’s adjuvant (CFA)-induced chronic inflammatory pain model

Female and male B6J and B6N mice were used (n = 6/sex/group) to study mechanical and thermal hypersensitivity after being administered CFA. Mice were injected in the dorsal surface of the right hind paw with 20 µL of CFA (10%). Mechanical hypersensitivity was measured via the von Frey test before injection and on day 3, 7, 14, 21, and 28 after CFA injection. On same days, paw edema was also evaluated. Thermal hypersensitivity was measured via the Hargreaves test on day 4, 10, 17, and 24 after CFA injection. The results shown pertain to the ipsilateral paw of each mouse.

### Chronic constrictive nerve injury (CCI)-induced neuropathic pain model

To evaluate possible differences in the development of CCI-induced neuropathic pain-like behavior, female and male B6J and B6N (n=6/sex/group) were tested in CCI neuropathic pain model. Mice were anesthetized using 3-4% isoflurane and maintained with 1-2% isoflurane in oxygen using a face mask and a vaporizer (VetEquip Inc, Pleasanton, CA). An incision was made just below the left hip bone, parallel to the sciatic nerve. The left common sciatic nerve was exposed at the level proximal to the sciatic trifurcation and a nerve segment 3-5 mm long was separated from surrounding connective tissue. Two loose ligatures with 5-0 silk suture were made around the nerve with a 1.0-1.5 mm interval between each of them. The wound was closed with suture thread. This procedure resulted in CCI of the ligated nerve and induced neuropathic insult. Animals were tested for changes in mechanical hypersensitivity before surgery and at days 3, 5, 7, 11, 14, 21, 28, and 35 post-surgery. The results shown pertain to the ipsilateral paw of each mouse.

### Mechanical hypersensitivity

Mechanical withdrawal thresholds were determined according to the method of Chaplan et al.^(24)^ with slight modifications^(25)^. Mice were placed in clear plastic cylinders (9 × 11 cm) with mesh metal flooring and allowed to acclimate for 15 min before testing. A series of calibrated von Frey filaments (Stoelting, Inc., Wood Dale, IL USA) with logarithmically incremental stiffness ranging from 2.83 to 4.56 units expressed Log 10 of [10 x force in (mg)] were applied to the paw using a modified up-down method^(25)^. Testing commenced with the 3.84 numbered filament for each mouse and continued up or down according to response. In the absence of a paw withdrawal response to the initially selected filament, a thicker filament corresponding to a stronger stimulus was presented. In the event of paw withdrawal, the next weaker stimulus was chosen. Each hair was presented perpendicularly against the paw, with sufficient force to cause slight bending, and held 2-3 s. The stimulation of the same intensity was applied two times to the hind paw at intervals of a few seconds. The mechanical threshold (paw withdrawal threshold, **PWT**) was expressed in g, indicating the force of the von Frey hair to which the animal reacted (paw withdrawn, licking, or shaking). All behavioral testing on animals was performed in a blinded manner. The data is expressed with the results of the ipsilateral paw of each mouse.

### Thermal hypersensitivity

Thermal withdrawal latencies were measured via the Hargreaves test by placing the mice in clear plastic cylinders (9 × 11 cm) with glass plate flooring and allowing them to acclimate for 15 min before testing. An infrared heat emitter was used to detect the thermal hypersensitivity at an intensity of 2.8. The infrared emitter was placed under the paw of the mouse and the paw withdrawal latency (**PWL**) was recorded for both the each hind paw of each mouse. An average of 2-3 measures of the paw withdrawal latency were taken for each hind paw. Results are shown for the ipsilateral paw of each mouse.

### Paw edema

The thickness of the formalin- or CFA-treated paws were measured both before and after injections on certain time points using a digital caliper (Traceable Calipers, Friendswood, TX USA). Data were recorded to the nearest ±0.01 mm and expressed as change in paw thickness (ΔPD = difference in the ipsilateral paw diameter before and after injection paw thickness).

### Hot plate test of thermal nociception

B6J (n = 24; 12 females, 12 males) and B6NJ (n = 18; 11 females, 7 males) parental substrains as well as B6J x B6NJ-F2 mice (N = 164) were tested on the 52.5°C hot plate assay^(26, 27)^. Mice were habituated to the testing room for at least 1 h. Mice were then placed in a Plexiglas cylinder (15 cm diameter; 33.0 cm tall) on a hot plate (IITC Life Science Inc., Woodland Hills, CA, USA) and the latency to lick the hind paw with a 60 s cut-off latency was recorded using a stopwatch. The B6J and B6NJ parental substrains were experimentally naïve at the time of nociceptive assessment. F2 mice were part of a larger historical dataset in which mice had a prior history of training in a conditioned place preference protocol as described^(28)^ that involved two training injections of either saline (SAL; i.p.; n=83) or the mu opioid receptor agonist oxycodone (OXY; 1.25 mg/kg, i.p.; n=81). During the four days prior to baseline hot plate assessment, F2 mice had continued to receive four daily injections of either SAL (i.p.) or OXY (20 mg/kg, i.p.). Twenty-four h after the fourth injection, mice were assessed for baseline hot plate sensitivity.

### DNA collection and genotyping in F2 mice

DNA was extracted from spleens and prepared for genotyping using a standard salting out protocol. Ninety SNP markers spaced approximately 30 Mb (approximately 15 cM) apart were genotyped using a custom-designed Fluidigm array (South San Francisco, CA USA). This level of coverage is sufficient for initial mapping in an F_2_ cross because of the low number of recombination events. We recently used a nearly identical marker panel for QTL mapping in the same cross^(19)^. We included an additional marker on chromosome 7 (rs31995355; 4 Mb) to improve QTL resolution. Markers were selected using high coverage sequence data for the C57BL/6NJ strain generated by the Welcome Trust Sanger Institute^(15, 29)^ and were validated using traditional Sanger sequencing. Genomic DNA was diluted to 100 ng/uL in low TE buffer (Teknova, Hollister, CA, USA) and genotyped with approximately 20% replication using the Fluidigm 96 × 96 SNPtype assay according to the manufacturer instructions. SNPs were called using the Fluidigm SNP Genotyping Analysis Software and SNPtype Normalization with the default 65% confidence threshold.

### RNA collection, library preparation, and sequencing for expression QTL (eQTL) analysis

Striatum punches were collected as described^(30)^ for RNA-seq from 23 F2 mice (all OXY-treated for historical reasons). Brains were rapidly removed and sectioned with a brain matrix to obtain a 3 mm thick section where a 2 mm diameter punch of the striatum was collected. Left and right striatum punches were pooled and immediately placed in RNAlater (Life Technologies, Grand Island, NY, USA) for 48 h prior to storage in a −80°C freezer. Total RNA was extracted using the RNeasy kit (Qiagen, Valencia, CA, USA) as described^(30)^. RNA was shipped to the University of Chicago Genomics Core Facility for cDNA library preparation using the Illumina TruSeq (oligo-dT; 100 bp paired-end reads). Libraries were prepared according to Illumina’s detailed instructions accompanying the TruSeq® Stranded mRNA LT Kit (Part# RS-122-2101). The purified cDNA was captured on an Illumina flow cell for cluster generation and sample libraries were sequenced at 23 samples per lane over 5 lanes (technical replicates) according to the manufacturer’s protocols on the Illumina HiSeq 4000 machine, yielding an average of 69.4 million reads per sample. FASTQ files were quality checked via FASTQC and possessed Phred quality scores > 30 (i.e. less than 0.1% sequencing error).

### Statistical Analysis

Behavioral data from the formalin test, CFA, and CCI were analyzed using the GraphPad Prism software, version 6.0 (GraphPad Software, Inc., La Jolla, CA) and expressed as the mean ± S.E.M. Statistical analysis was conducted using analysis of variance test (**ANOVA**) and followed by a post hoc test. Before ANOVA, the data were first assessed for the normality of the residuals and equal variance. Variances were similar between groups and were assessed using either the F-test or the Brown–Forsythe test and the Bartlett’s test. All data passed these tests. A two-way ANOVA followed by the Tukey’s post hoc correction was used in formalin test. In addition, an unpaired student *t* test was used to compare paw thickness in the formalin test. To test the mechanical and thermal sensitivity, repeated measures (**RM**) two-way ANOVA was used with Sidak (to compare substrain effects) and Dunnett’s (to compare time’s effect according to baseline value) post hoc correction. A p value of less than 0.05 was considered significant.

QTL analysis was performed in F_2_ mice using the R package R/qtl as previously described^(19, 31, 32)^. The scanone function was used to calculate LOD scores. Permutation analysis (perm = 1,000) was used to establish the significance threshold for each quantitative trait (p<0.05). The marker position (cM) was estimated using the sex-averaged position using the Mouse Map Converter (http://cgd.jax.org/mousemapconverter)^(33)^. The converter tool was also used to estimate the Mb position of the QTL peak and the Bayes credible interval that defined the QTL region. The 90 polymorphic markers used for genetic mapping and the reported variants that underlie the hot plate QTL interval were retrieved from the Sanger database (http://www.sanger.ac.uk/). Power analysis was used to inform the F2 sample size for QTL analysis and was conducted using GPower (http://www.gpower.hhu.de/en.html). Using the means, standard deviations, and unpaired two-tailed t-tests, we calculated the effect size (Cohen’s d) of the parental B6 substrain difference in hot plate latency. We then used this effect size and set the alpha level to 0.05 and power level to 95% to estimate the required sample size for each homozygous genotype at a given locus.

For expression QTL mapping, we aligned FastQ files to the reference genome (mm38) via TopHat^(34)^ using the mm38 build and Ensembl Sequence and genome annotation. We used *featureCounts* to count and align reads. For *cis*-eQTL analysis, we used the same 90 SNPs that were used in behavioral QTL analysis. We removed lowly expressed exons that did not possess at least 10 reads total across all 115 count files. Because of the low resolution of QTL mapping in an F2 cross, we liberally defined a gene with a *cis*-eQTL as any gene possessing a genome-wide significant association between expression and a polymorphic marker that was within 70 Mb of a SNP (the largest distance between any two SNPs from the 90-SNP panel). The genes reported within the chromosome 7 interval all showed their most significant association with gene expression at the peak SNP associated with hot plate sensitivity (rs3148686; 30.31 Mb). Analysis was conducted using *limma* with default TMM normalization and VOOM transformation^(35)^. A linear model was employed whereby sample replicates were treated as a repeated measure. The duplicateCorrelation() function was used to estimate within-sample correlation, which was then included in the lmFit() function. An ANOVA test was conducted for gene expression that included Sex as a covariate and Genotype as a fixed effect. Gene-level tests were conducted using the likelihood Ratio test. A false discovery rate (**FDR**) of 5% was employed as the cut-off for statistical significance^(36)^.

## RESULTS

### Formalin-induced inflammatory nociceptive behaviors in B6J and B6N substrains

As seen in **Figure 1A**, we evaluated the paw licking responses of B6J and B6N mice in the formalin test at 2.5% concentration. Two-way ANOVA revealed significant effects on time spent in paw licking time in terms of substrain [F_strain_(1,44)=12.91, p<0.001], phase [F_phase_(1,44)=160.1, p<0.001] and interaction [F_interaction_(1,44)=4.68, p<0.05]. Although no significant difference was found in phase I behaviors between B6J and B6N mice (p>0.05, **Figure 1A**), a significant increase in paw licking time in B6J mice compared to B6N mice was found in the phase II (p<0.001, **Figure 1A**). Moreover, the degree of paw edema (thickness) in B6J mice was significantly greater than B6J mice at 1 h post-formalin injection [t(22)=4.163, p<0.001, **Figure 1B**].

**Figure 1.**
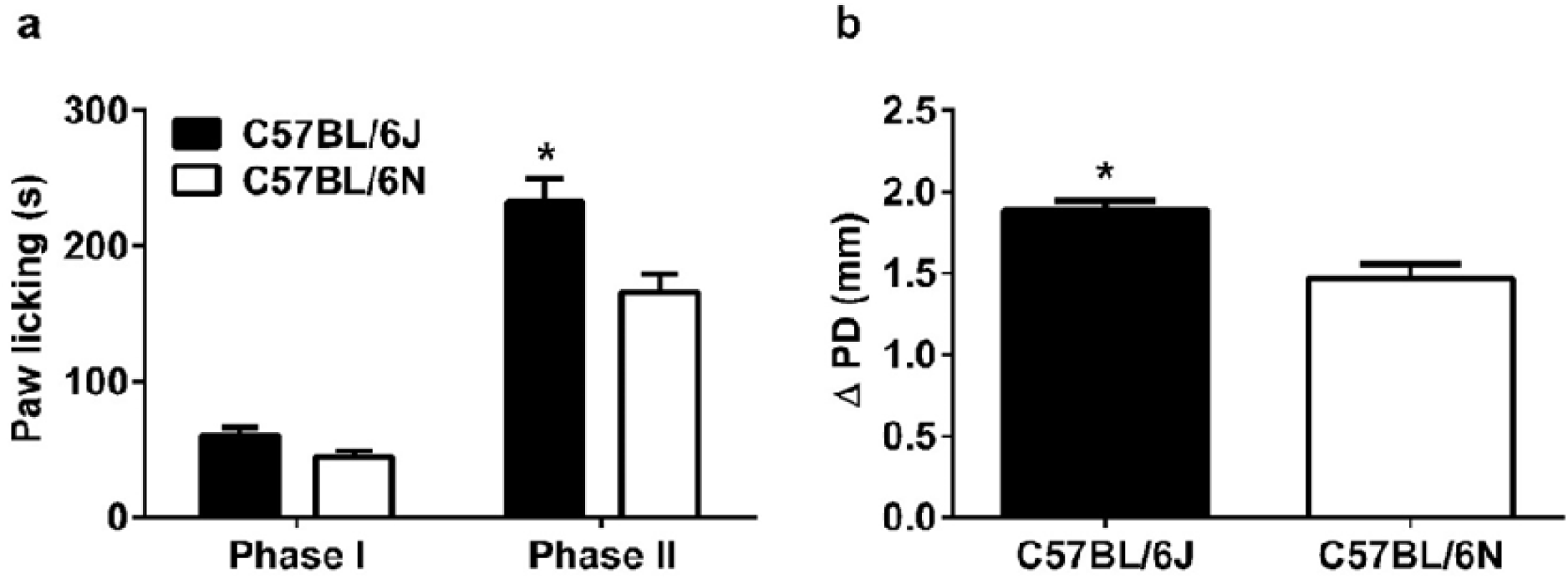
Formalin-induced paw licking behavior and paw edema in B6J and B6N mice. The paw licking response after injection of **(A)** 2.5% formalin concentration into the right paw of both B6J and B6N mice. Changes in paw edema **(B)**, as measured by the difference in the ipsilateral paw diameter before and after injection (ΔPD), in B6J & B6N mice 1 hour after injection of formalin. Data are expressed as the mean ± S.E.M. of 6 mice/per sex/per group. *p<0.05 significantly different from B6J mice.

**Supplementary Figure 1** shows the effect of formalin on paw licking and edema in females and males. A reduction on time spent in paw licking was found in male B6N mice {[F_substrain_(1,20)=9.83, p<0.01], phase [F_phase_(1,20)=67.36, p<0.001] and interaction [F_interaction_(1,20)=3.78, p=0.66]} (**Supplementary Figure 1A**). Consistent with the paw licking behavior results, paw diameter was lower in B6N male mice compared to B6J males [t=4.163, df=22, p<0.001, **Supplementary Figure 1B**]. However, there was no significant difference on paw licking {[F_substrain_(1,20)=3.08, p=0.094], phase [F_phase_(1,20)=88.28, p<0.001] and interaction [F_interaction_(1,20)=0.9843, p=0.33]} (**Supplementary Figure 1C**) and paw edema between B6J female and B6N female mice [t(22)=4.16, p<0.001, **Supplementary Figure 1D**].

### CFA-induced mechanical and thermal hypersensitivity in B6J and B6N substrains

Mice were given an injection of CFA (10%) and tested for mechanical and thermal sensitivity at several time points (days) before and after post-CFA injection. In addition, paw edema was evaluated. For mechanical responses, two-way ANOVA showed significant effects of time [F_time_(6,66)=22.57, p<0.001], but not for substrain [F_substrain_(1,11)=0.4384, p=0.52] or interaction [F_interaction_(6,66)=0.33, p=0.92] (**Figure 2A**). Prior to CFA injection, B6J and B6N mice did not differ in mechanical baseline values (p>0.05, **Figure 2C**), similar to observations from a recent study^(23)^. The CFA injection induced a decrease in mechanical thresholds which indicates an induction of mechanical hypersensitivity by CFA in both substrains (**Figure 2A**). Significant mechanical hypersensitivity started by day 3 post-CFA injection (p<0.05) and continued until day 21 (p>0.05, **Figure 2A**). Furthermore, we tested thermal paw withdrawal latencies on different days than paw withdrawal threshold testing. Two-way ANOVA showed significant effects for thermal responses for time [F_time_(4,44)=41.05, p<0.001], but not for substrain [F_substrain_(1,11)=3.25, p=0.099] and interaction [F_interaction_(4,44)=0.82, p=0.52] (**Figure 2B**). Prior to CFA injection, baseline values for paw withdrawal latencies were similar between B6J and B6N mice (p>0.05, **Figure 2B**). CFA injection resulted in a significant reduction on paw withdrawal latencies which indicates the formation of thermal hypersensitivity (p<0.05). The thermal hypersensitivity was observed on days 11 and 17 after CFA injection (p<0.05); mice were recovered from thermal hypersensitivity by day 24 (p>0.05, **Figure 2B**). CFA-induced paw edema did not differ between B6J and B6N mice [F_time_(5,55)=35.14, p<0.001; F_substrain_(1,11)=0.66, p=0.4338; F_interaction_(5,55)=1.72, p=0.15; **Figure 2C**].

**Figure 2.**
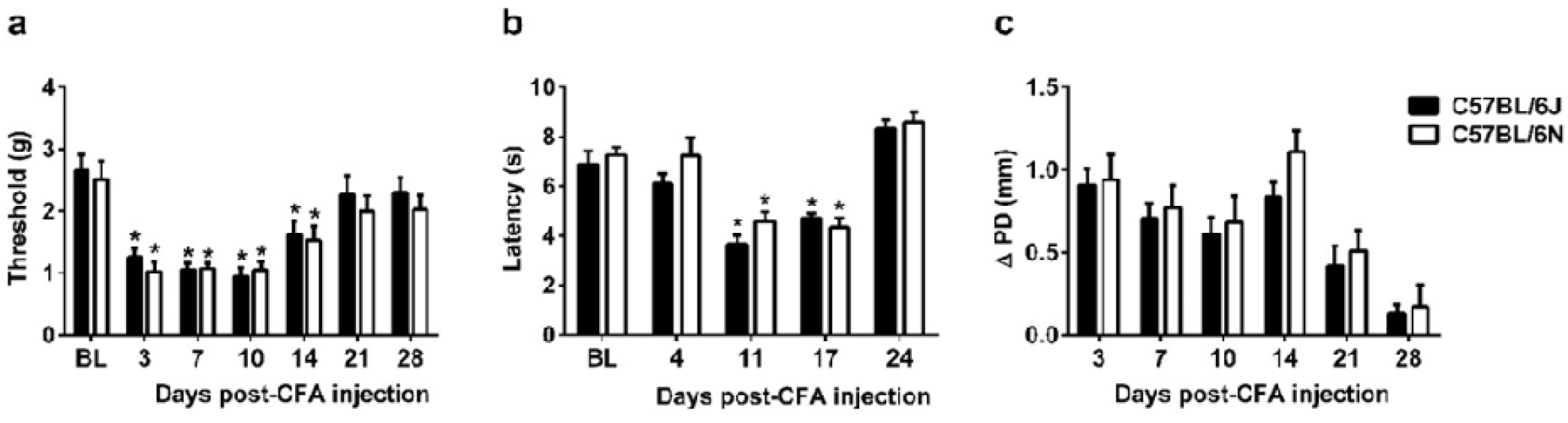
CFA-induced mechanical & thermal hypersensitivity and paw edema in B6J and B6N mice. Differences in **(A)** mechanical paw withdrawal thresholds and **(B)** paw withdrawal latencies (Δ PWL=contralateral–ipsilateral hindpaw latencies) in B6J and B6N mice at different times after injection of complete Freund’s adjuvant (CFA, 10 % solution/20 µl). Degree of edema **(C)**, as measured by the difference in the ipsilateral paw diameter before and after injection (ΔPD) in B6J and B6N mice. Data are expressed as the mean ± S.E.M. of 6 mice/per sex/per group. *p<0.05 significantly different from the value of baseline (BL)

**Supplementary Figure 2** shows a comparison of CFA-induced mechanical and thermal hypersensitivity in B6J and B6N substrains for females and males. Two-way ANOVA showed no significant substrain differences in mechanical withdrawal thresholds [F_time_(6,30)=14.11, p<0.001; F_substrain_(1,5)=6.155, p=0.056; F_interaction_(6,30)=0.18, p=0.98; **Supplementary Figure 2A**] or in thermal withdrawal latencies [F_time_(4,20)=23.17, p<0.001; F_substrain_(1,5)=0.91, p=0.39; F_interaction_(4,20)=0.76, p=0.56; **Supplementary Figure 2B**]. Similarly, there was no significant substrain differences in mechanical withdrawal thresholds [F_time_(6,30)=10.58, p<0.001; F_substrain_(1,5)=1.11, p=0.34; F_interaction_(6,30)=0.26, p=0.95; **Supplementary Figure 2C**] or in thermal withdrawal latencies [F_time_(4,20)=16.28, p<0.001; F_strain_(1,5)=2.632, p=0.17; F_interaction_(4,20)=4.41, p<0.05; **Supplementary Figure 2D**] between female B6J and B6N mice.

### CCI-induced mechanical hypersensitivity in B6J and B6N substrains

The development of CCI-induced mechanical hypersensitivity in von Frey test, was compared in B6J and B6N mice. In examining mechanical thresholds over time, two-way ANOVA indicated a main effect of time [F_time_(8,88)=164.4, p<0.001], but not substrain [F_strain_(1,11)=0.74, p=0.41] or interaction [F_interaction_(8,88)=1.29, p=0.26] (**Figure 3**). Prior to CCI surgery, B6J and B6N mice did not differ in mechanical baseline values (p>0.05, **Figure 3**). A reduction in mechanical paw withdrawal thresholds was observed at 3 days post-surgery and continued over 35 days (p<0.05, **Figure 3**). The mechanical hypersensitivity was similar between substrains. Moreover, male [F_time_(8,40)=62.87, p<0.001; F_substrain_(1,5)=1.401, p=0.2898; F_interaction_(8,40)=1.347, p=0.2491; Supplementary Figure 3a] and female [F_time_(8,40)=106.5, p<0.001; F_substrain_(1,5)=0.0696, p=0.8024; F_interaction_(8,40)=0.4246, p=0.8993; **Supplementary Figure 3B**] showed similar mechanical hypersensitivity in B6J versus B6N substrains.

**Figure 3.**
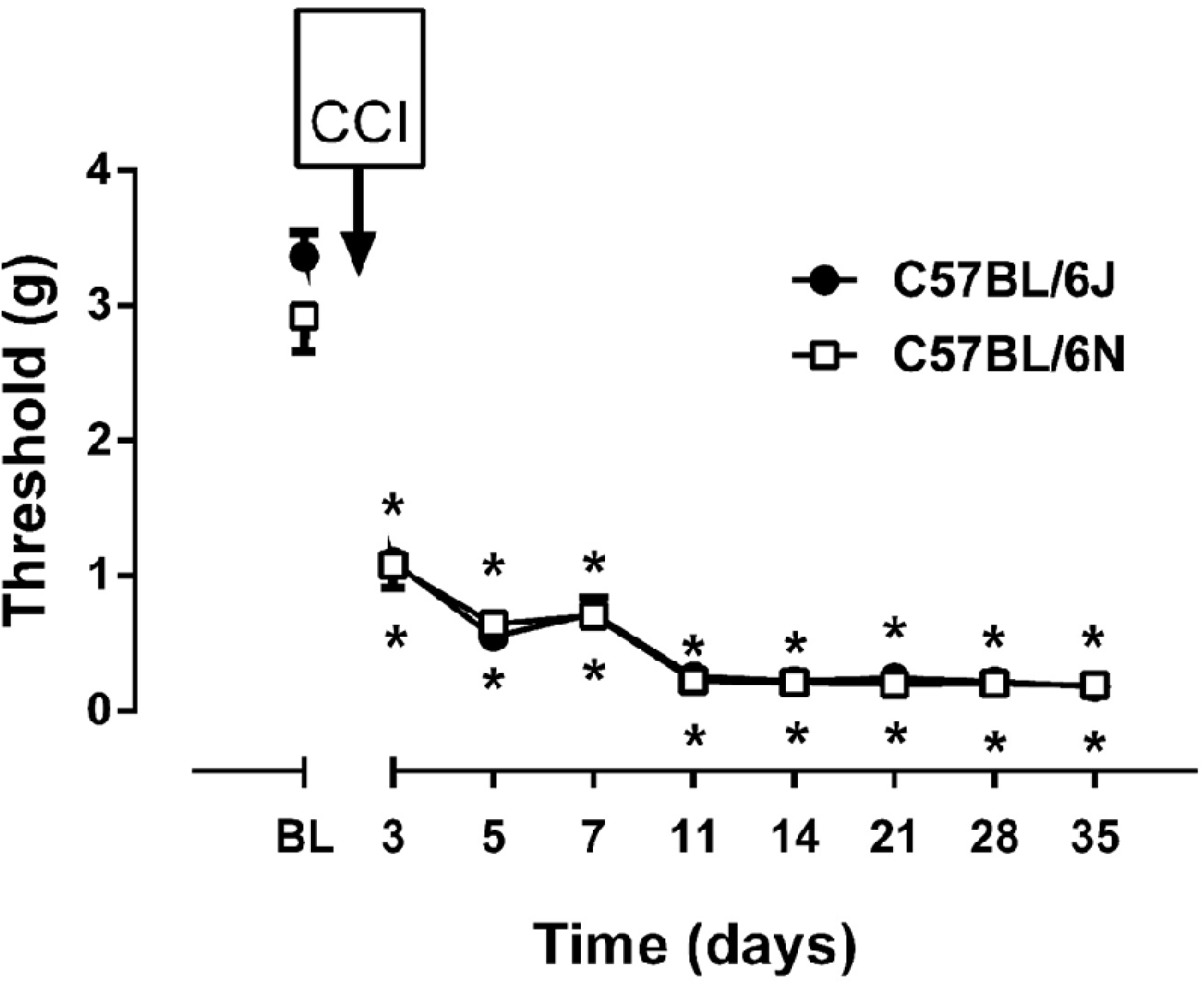
CCI-induced mechanical and thermal hypersensitivity in B6J and B6N mice. Differences in mechanical paw withdrawal thresholds in B6J and B6N mice at different days after chronic constrictive nerve injury (CCI) operation. Data are expressed as the mean ± S.E.M. of 6 mice/per sex/per group. *p<0.05 significantly different from the value of baseline (BL).

### Increased sensitivity to acute, thermal nociception in the hot plate test in B6J versus B6NJ substrains

We and others previously reported an increase in acute thermal nociceptive sensitivity (i.e., decreased baseline latencies) in the B6J substrain versus the B6N (Crl) substrain^(21, 22)^. Here, we extended this finding in comparison with another genetically very similar B6N “sub”-substrain^(37)^ for which whole genome sequence information is available^(15, 16, 29)^ – the B6NJ substrain from JAX [**Fig.4A**: t(40)=3.59; p=8.9 × 10^−4^]. When breaking down the data by females and males, the difference was significant with females [t(21)=2.08; p=0.0075; n=12 B6J, n=12 B6NJ) but did not reach the p<0.05 cut-off (two-tailed) for significance in males [t(17)=2.11; p=0.078], likely due the smaller sample size; n=12 B6J, n=7 B6NJ; **Supplementary Figure 4A**]. However, based on our historical findings and others in males^(21, 22)^ a one-tailed test is justified for the males and yields the expected significant decrease in B6J versus B6NJ males (t17=1.74; p=0.039) (**Supplementary Figure 4A**).

**Figure 4.**
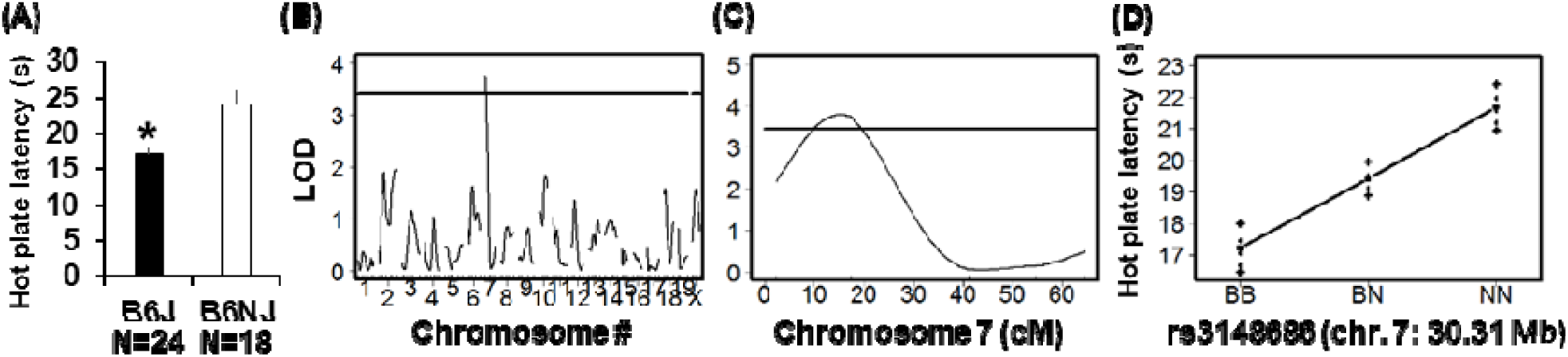
A major QTL on chromosome 7 underlies B6 substrain differences in acute, thermal nociception in the 52.5°C hot plate assay. **(A)**: We replicated our previous observation of a significant increase in sensitivity to acute thermal nociception (decreased in latency to lick the hindpaw) in the parental B6J substrain versus the B6NJ substrain^(21)^ (t_40_ = 3.59; p = 8.9 × 10^−4^). **(B):** Genome-wide significant QTL on chromosome 7 was identified from 164 B6J x B6NJ-F2 mice [LOD = 3.81; peak = 17 cM (6 Mb); peak marker (rs3148686) = 30 Mb; Bayes C.I.: 8.74 Mb – 36.50 Mb). **(C):** QTL plot for chromosome 7 is shown. Horizontal line for panels B and C denotes the significance threshold (1000 permutations). **(D):** Effect plot at the peak-associated marker illustrates the decreased hot plate latency associated with the B6J allele (B) and an additive effect with inheritance of one copy versus two copies of the B6NJ allele (N). BB = homozygous for B6J allele; BN = heterozygous; NN = homozygous for B6NJ allele. Data for panels A and D are expressed as the mean ± SEM.

### Identification of a major QTL on chromosome 7 underlying enhanced acute thermal nociceptive sensitivity in B6J versus B6NJ substrains

Power analysis using the effect size estimated from the hot plate latencies of the parental strains (Cohen’s d=1.07) indicated that a sample size of n=24 homozygous genotypes was required to achieve 95% power (p<0.05). We employed a sample size of n=164 which yielded n = 39 B6J and n=42 B6NJ genotypes at the peak-associated marker and is within range of the predicted n=41 per homozygous genotype based on a 1:2:1 Mendelian ratio. Using this sample size, we achieved greater than 99% power to detect a significant difference of the predicted effect size of d = 1.07 (p < 0.05). If we adjust the alpha level to 0.00056 to account for the 90 statistical tests (0.05/90), we still achieve 88% power (p=0.00056) to detect an effect of d=1.07, given our homozygous genotype sample sizes of n=39 homozygous B6J and n=42 homozygous B6NJ at the peak QTL marker.

A robust phenotypic difference in hot plate sensitivity combined with availability of polymorphic markers and high throughput genotyping tools equipped us with the ability to accomplish our longstanding goal of conducting a genome-wide QTL study of baseline thermal nociception in a cross between B6 substrains^(17)^. A list of the 90 polymorphic markers used in QTL analysis is provided in **Supplementary Table 1**. We identified a single genome-wide locus on chromosome 7 underlying B6 substrain differences in baseline nociception as measured via the hot plate assay [LOD = 3.81; QTL peak = 14 cM (26.14 Mb); Bayes Credible Interval = 5.30 cM – 22.30 cM (8.74 Mb–36.50 Mb); **Fig. 4B,C**]. The peak marker, rs3148686, is located at 30.31 Mb. Importantly, the QTL signal was driven by allelic differences in the predicted phenotypic direction (**Fig. 4D** versus **Fig.4A; Supplementary Figure 4A**). Mice with a prior history of OXY versus SAL injections showed similarly trending QTLs and genotypic pattern of hot plate latencies or QTL detection (**Supplementary Figure 5**). Furthermore, inclusion of Prior Treatment as a covariate into the QTL model still yielded a significant LOD score (LOD with covariate = 3.72, p<0.01). The effect size of this locus in comparing the homozygous phenotypes was d = 0.87 and thus, we achieved 70% power (p<0.00056). To summarize, we identified a major genetic locus underling enhanced acute thermal pain sensitivity in B6J versus B6N mice. Surprisingly, when we conducted QTL analysis separately in females and males, the chromosome 7 QTL was only significant in males (**Supplementary Figure 5B-F**).

### Striatal *cis*-expression QTLs within chromosome 7 QTL for hot plate sensitivity

The RNA-seq gene expression dataset has been uploaded to Gene Expression Omnibus (GEO) and is available to the reviewer at the following link: https://www.ncbi.nlm.nih.gov/geo/query/acc.cgi?acc=GSE119719. The reviewer access token is “azmtaeseplipbyv”

Complementary eQTL analysis can identify functionally relevant candidate genes, providing a link between genotype, gene expression, and behavior^(38)^. We took advantage of a historical dataset that we generated from striatal brain tissue collected from 23 OXY-treated F2 mice in order to document genes within the chromosome 7 QTL that also possess *cis*-eQTLs. OXY-treated mice also show variation in acute thermal pain sensitivity that maps to the same chromosome 7 hot plate QTL (**Supplementary Figure 5**); thus, gene expression in OXY-treated mice is still relevant to the behavioral QTL.

The genes we identified were located within 1 Mb to 17 Mb distance of the peak-associated marker for hot plate latency. We identified 15 genes whose expression was modulated by *cis*-eQTLs (FDR < 0.05) (**Table 1**). Eight of these genes contain known variants (*Ryr1, Rps5, Cox6b1, Pou2f2, Clip3, Sirt2, Actn4*, and *Ltbp4*)^(15)^ (**Supplementary Table 2**) and nine of these genes have been implicated in either pain and/or inflammation (*Ryr1, Cyp2a5, Rps5, Gys1, Pou2f2, Clip3, Sirt2, Actn4*, and *Ltbp4*). Based on the distance from the peak-associated SNP (1 Mb), the strength of the association (FDR = 1.5 × 10^−5^), the number of variants overlapping the cognate gene (9 noncoding SNPs), and representation in the pain literature, we nominated the ryanodine receptor 1 (*Ryr1*; 29 Mb) as a high priority candidate gene underlying enhanced hot plate sensitivity. The nociception-enhancing B6J allele was associated with a 1.25-fold decrease in *Ryr1* expression (or i.e., the B6NJ allele showed an increase in expression; **Table 1**). Only two other genes besides *Ryr1* showed a larger fold-change in expression, including two cytochrome P450 genes - *Cyp2a5* and *Cyp2g1* in which the B6J allele showed a 3.6- to 3.8-fold decrease in expression relative to the B6NJ allele (or i.e., B6NJ allele showed an increase in expression; **Table 1**). *Cyp2a5* (26.84 Mb) and *Cyp2g1* (26.81 Mb) are located very near peak marker associated with the chromosome 7 QTL for hot plate latency that was inferred from interval mapping (26 Mb).

**Table 1.**
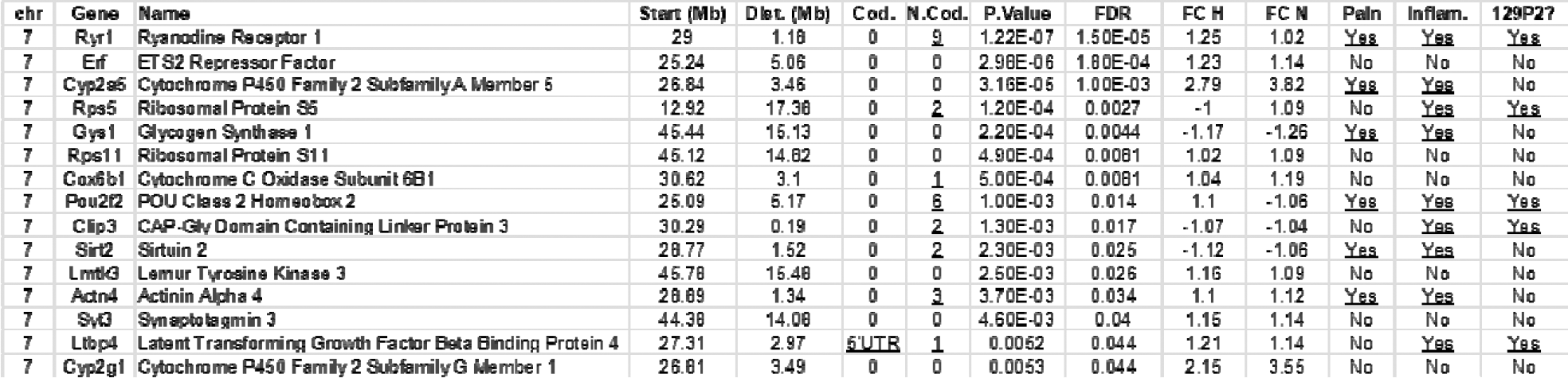
Genes within the hot plate QTL on chromosome 7 that have a *cis*-eQTL associated with rs3148686. “Dist.(Mb)” = distance of gene from the SNP rs3148686 (30.31 Mb), the SNP nearest the QTL for hot plate sensitivity. “Cod.” Refers to coding polymorphisms. “N.Cod.” refers to non-coding polymorphisms. FDR = false discovery rate. FC H = fold-change in gene expression in mice heterozygous (H) for the rs3148686 marker relative to mice homozygous for the B6J (B) allele.FC N = fold-change in gene expression in mice homozygous for the B6NJ allele (N) relative to mice homozygous for the B allele. “Pain” refers to a pain literature search (“gene” and “pain” or “nociception”; pubmed). “Inflam.” Refers to an inflammation literature search (“gene” and “inflammation” or “inflammatory”; pubmed). “129P2?” refers to whether or not the gene contains a variant shared between the nociception-resistant B6NJ strain and the 129P2 strain.

## DISCUSSION

We expanded the catalog of behavioral differences in nociceptive sensitivity between B6 substrains^(21, 22)^ and identified enhanced inflammatory nociception in the formalin test in the B6J versus B6N substrain (**Fig.1**) in the absence of any behavioral differences in the more chronic CFA inflammatory pain model (**Fig.2**) or in the CCI neuropathic pain model (**Fig.3**). Thus, there is selectivity of enhanced nociceptive sensitivity in the B6J strain with regard to both the nociceptive modality as well as the particular noxious stimulus within a nociceptive modality (inflammatory). In the second part of the study, we replicated the enhanced sensitivity to acute thermal nociception as measured via the hot plate assay in B6J versus B6N substrains from JAX (**Fig.4A**) and mapped a major QTL underlying parental substrain differences on chromosome 7 (**Fig.4B,C**) that mirrored the parental substrain difference (**Fig.4D**). Finally, using a historical striatal eQTL dataset, we identified 15 genes possessing eQTLs within 1 to 17 Mb of the peak-associated marker, providing functional support for their candidacy.

The B6J substrain showed a more pronounced and statistically significant inflammatory nociceptive response during the late phase of the formalin test (**Figure 1A**). The early phase of nociceptive behavior (e.g., 0-5 min) is thought to model acute chemical pain and activation of c-fibers whereas the late phase (e.g., 20-45 min) is thought to model an inflammatory response and functional changes in the dorsal horn of the spinal cord^(39)^. The larger increase in paw diameter following the formalin test in the B6J substrain further supports an increased inflammatory response (**Figure 1B**). Male B6J mice showed a more robust increase in nociceptive behaviors relative to their B6N male counterparts compared to females during the late phase (**Supplementary Figure 1**); however both female and male B6J mice showed a significant increase in paw diameter relative to their B6N counterparts. The former observation could be due to chance sampling error and/or to the effects of sex steroids on formalin-induced nociceptive behaviors^(40-43)^ that obfuscate detection of genetic differences.

Interestingly, the genome-wide significant QTL on chromosome 7 that we identified for hot plate thermal nociception was only significant for male mice (**Supplementary Figure 4B-D**). The reason for this observation is not entirely clear. Our previously published results and others that showed enhanced thermal nociception in B6J versus B6N mice were collected in males only^(21, 22)^. However, in the present study, our parental substrain results were generated from both females and males and when considering only females, we also observed a significant decrease in hot plate latency in B6J versus B6NJ (**Supplementary Figure 4A**). Thus, the lack of detection of a QTL from female mice cannot be explained by a lack of *a priori* evidence in females or, e.g., by the fact that we used a different B6N substrain compared to our previous study^(21)^. Given the trending effect plots in for the females in the QTL study (**Supplementary Figure 4F**), one could speculate that with a larger sample size we may have detected a significant QTL. Additionally, we did not monitor the estrus cycle which is known to influence nociception, e.g., hot plate nociception^(44)^ and tail flick nociception^(45)^ and chronic/persistent pain^(46)^ and thus, variation in the phase of the estrus cycle could affect nociceptive sensitivity and obscure the effect of genotype. The sex of the experimenter is known to influence nociceptive latencies in mice^(47)^; however, in this case the experimenters for phenotyping the parental substrains (S.L.K.) and the F2 mice (L.R.G.) were both females. Finally, it is possible that a larger sample size might have revealed a genome-wide, female-specific locus.

The QTL on chromosome 7 overlaps with a QTL previously identified for acute thermal nociceptive sensitivity in the 49°C tail withdrawal assay in an F2 cross between B6J and 129P3 strains^(48)^. The QTL peak for tail withdrawal sensitivity was more distally located at 33 cM (C.I.: 24-38 cM) but was also associated with enhanced pain sensitivity in mice possessing the B6J allele. To our knowledge, the 129P3 strain has not been whole genome-sequenced and thus, a complete catalog of variants is lacking. However, whole genome sequence data is available for the 129P2/OlaHsd substrain (https://www.sanger.ac.uk/), one of the founder 129 strains^(49)^ that is genetically similar to 129P3. Interestingly, C57BL/6NJ and 129P2 strains share variants within many of the genes listed in **Table 1** that possess *cis*- eQTLs, including *Ryr1, Rps5, Pou2f2, Clip3*, and *Ltbp4*^(15, 16)^. Thus, if a private variant in B6J underlies enhanced sensitivity to acute thermal nociception and if B6NJ and 129P2 strains (and by further extension, the 129P3 strain) share the alternate allele via identity by descent^(20)^, then eQTL-containing genes that possess these SNPs are high priority candidates to pursue via gene targeting.

Interestingly, the chromosome 7 QTL we identified for hot plate nociception is localized to a nearly identical region reported for fluid intake of the nociceptive compound capsaicin (activates TRPV1) in an F2 cross between C57BL/6J and the wild-derived KJR strain^(50)^. The QTL was localized to approximately 19 cM (peak marker = D7Mit155; 31 Mb) which is nearly identical to the location for the hot plate QTL (peak marker = 30 Mb). The B6J allele was associated with decreased fluid intake of capsaicin compared to the KJR allele which is consistent with enhanced capsaicin-induced oral nociceptive sensation and is consistent with the overall theme of enhanced B6J-mediated nociception at this locus (**Figure 4**)^(48)^. Interestingly, a large panel of inbred strains tested alongside the C57BL/6 and KJR strains showed a similar pattern of strain differences in nociceptive sensitivity to capsaicin and hot plate^(51)^, indicating a genetic correlation and thus, a shared genetic basis between noxious chemical and thermal stimuli that generate heat sensation in their strain panel.

We examined gene expression in the striatum since this data was already available from a historical dataset. Arguably, because the nociceptive response in the hot plate assay is a supraspinally controlled, thermal acute response^(27)^, the relevant tissue for examining gene expression as it relates to thermal nociception could be a different tissue such as the dorsal root ganglia, the dorsal horn of the spinal cord, brainstem nuclei, midbrain periaqueductal gray, primary sensory cortex, or any of one of several limbic structures involve in supraspinal modulation of pain. In relation to the ventral striatum, calcitonin gene-related peptide (CGRP) which is an established pain neuropeptide that has been genetically mapped using inbred mouse strains^(12)^, induces an increase in hindpaw withdrawal latency in the hot plate when injected into the nucleus accumbens of rats (antinociception) and intra-accumbens injection of a CGRP antagonist induces a decrease in hot plate latency (hyperalgesia)^(52)^. Furthermore, activation of cyclic adenosine monophosphate response element-binding protein (CREB) in the nucleus accumbens modulates the hot plate response^(53)^. Thus, striatal tissue is potentially a relevant pain tissue^(54)^ as well as mesolimbic dopamine signaling within this brain region^(55)^ to understanding gene expression as it relates to genetic variation and nociception. It should also be noted that eQTLs from brain tissue are frequently expressed in multiple brain regions^(56)^. Thus, most of the genes we have identified will likely possess eQTLs in other CNS tissues that are relevant to thermal nociception.

*Ryr1* showed the strongest eQTL association, it possesses multiple polymorphisms, and is involved in calcium signaling in pain and inflammation. *Ryr1* codes for the ryanodine 1 receptor, a calcium release channel in the sarcoplasmic reticulum that is also associated with the dihydropyridine receptor, or i.e., L-type calcium channels. A large literature documents the involvement of L-type calcium channels in pain. Administration in male Swiss albino mice of ryanodine (i.c.v.), a Ryr antagonist dose-dependently reduced hot plate latencies whereas administration of 4-Cmc, a Ryr agonist, dose-dependently increased hot plate latencies^(57)^, providing direct pharmacological evidence for an involvement of central, supraspinal Ryr receptors and calcium release in acute thermal nociception. The reduced hot plate latency in response to the Ryr agonist is in line with the reduced hot plate latency in response that is associated with both reduced *Ryr1* expression and with the B6J allele (**Table 1**). Furthermore, administration of ryanodine and caffeine (Ryr agonist) in thalamocortical nuclei of B6 × 129 F1 mice produced similar bidirectional effects on both formalin-induced nociceptive behaviors during the late phase and acetic acid-induced writhing^(58)^, suggesting the possibility that differential Ryr1 expression could exert pleiotropic effects on both thermal and inflammatory nociception. On the other hand, another study showed that administration of verapamil (L-type blocker) in the dorsal horn in Sprague Dawley rats had no effect on formalin-induced nociception^(59)^.

In considering the fold-change in gene expression, it is striking that two cytochrome P450 genes, *Cyp2a5* (26.84 Mb) and *Cyp2g1* (26.81 Mb) are located squarely on top of the QTL peak for hot plate sensitivity and showed an extraordinary 3.8-fold and 3.6-fold change in transcription at the peak-associated hot plate marker, rs3148686 (**Table 1**). Notably, a nearby 112 kb structural variant (chromosome 7: 26.85-26.96 Mb) that is documented as an insertion with the B6J allele^(29)^ and as a deletion of the B6NJ allele^(15, 16, 29)^ (http://www.sanger.ac.uk/) is located just distally to *Cyp2g1* (26.81 Mb) and *Cyp2a5* (26.84 Mb). This structural variant includes only one protein-coding gene, *Cyp2a22* but is flanked by a cluster of CYP genes, including not only *Cyp2a5*, and *Cyp2g1* (**Table 1**) but also *Cyp2b23, Cyp2b19, Cyp2a12, Cyp2f2*, and *Cyp2t4*. The B6NJ deletion (or i.e., the B6J insertion^(29)^) could contain DNA sequences that normally serve to regulate *Cyp2a5* and *Cyp2g1* transcription or it could induce compensatory increases in *Cyp2a5* and *Cyp2g1* transcription through a separate mechanism.

To summarize, we identified robust B6 substrain differences in pain sensitivity in an additional nociceptive modality whereby the B6J strain was more sensitive to formalin-induced nociceptive behavior. These results support the previous conclusion by Mogil and colleagues decades ago that the B6J strain is not the ideal strain to study the molecular basis of nociceptive phenotypes as it is hypersensitive to multiple nociceptive modalities^(4, 60)^, yet a majority of molecular genetic studies continue to only utilize mice on a B6J background. A recent comprehensive study of F1 crosses between 30 inbred strains highlights the profound impact of epistatic modifier alleles in determining the magnitude and direction of knockout alleles on complex behavioral traits and the importance of studying mutations on multiple genetic backgrounds^(61)^. A limitation of the current study is that we only examined behaviors that model the sensory aspect of pain. An important question is whether or not there are B6 substrain differences in behaviors that model the motivational-affective and cognitive-evaluative components of pain. Future, additional genetic crosses between B6 substrains will determine the degree to which these different models of pain are genetically shared versus dissociable and rapid fine mapping strategies^(20)^ combined with gene editing will eventually identify the causal genetic factors.

## ACKNOWLEDGMENTS

R01CA221260 (M.I.D. and C.D.B.), R01DA039168 (C.D.B.), R21DA038738 (C.D.B.), T32GM008541 (L.R.G.)

## SUPPLEMENTARY INFORMATION

**Supplementary Table 1. can be found near the end of this document on p.8-p.9.**

**Supplementary Table 2. can be found at the end of this document on p.10**

**Supplementary Table 1.**
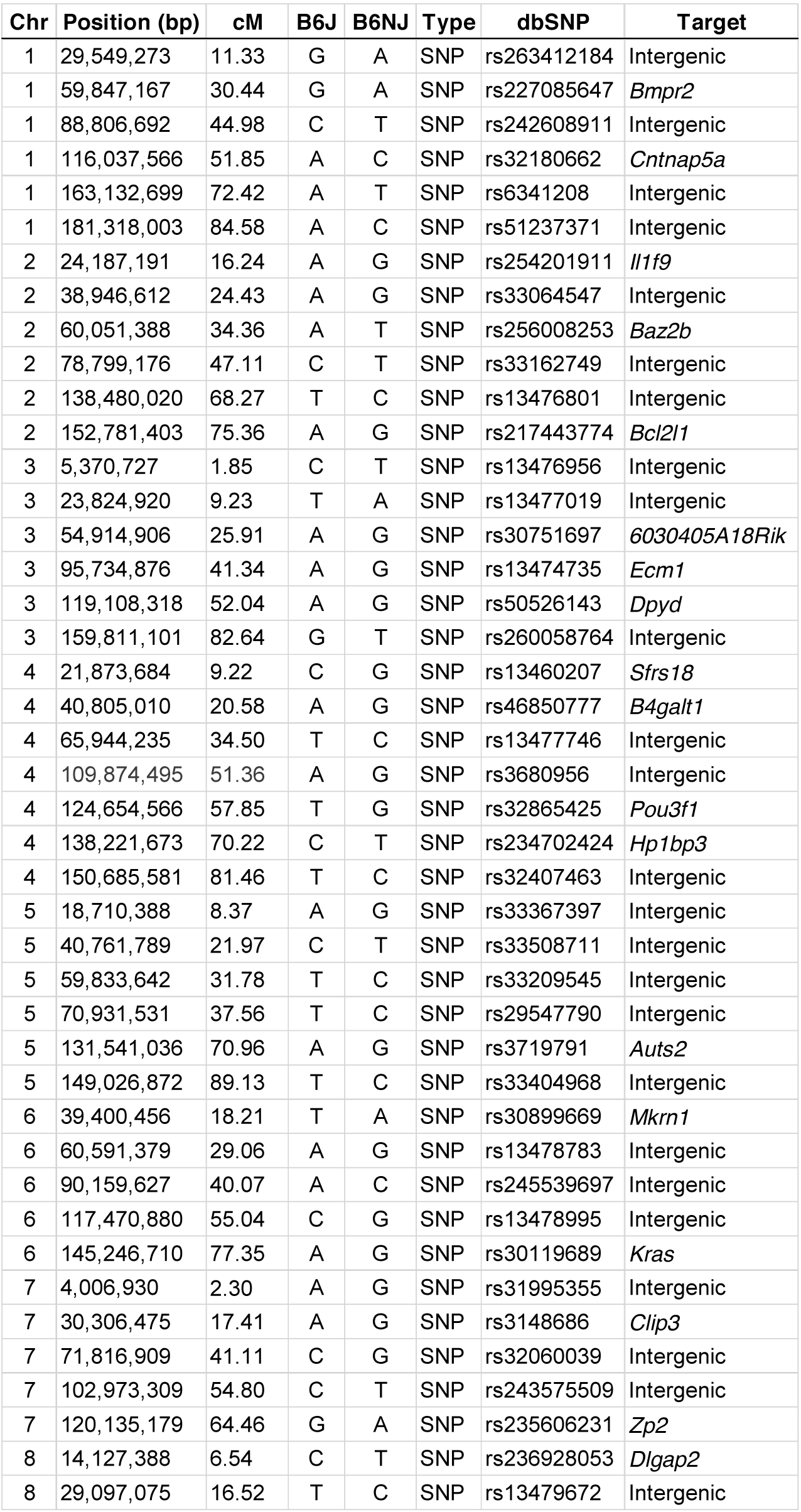
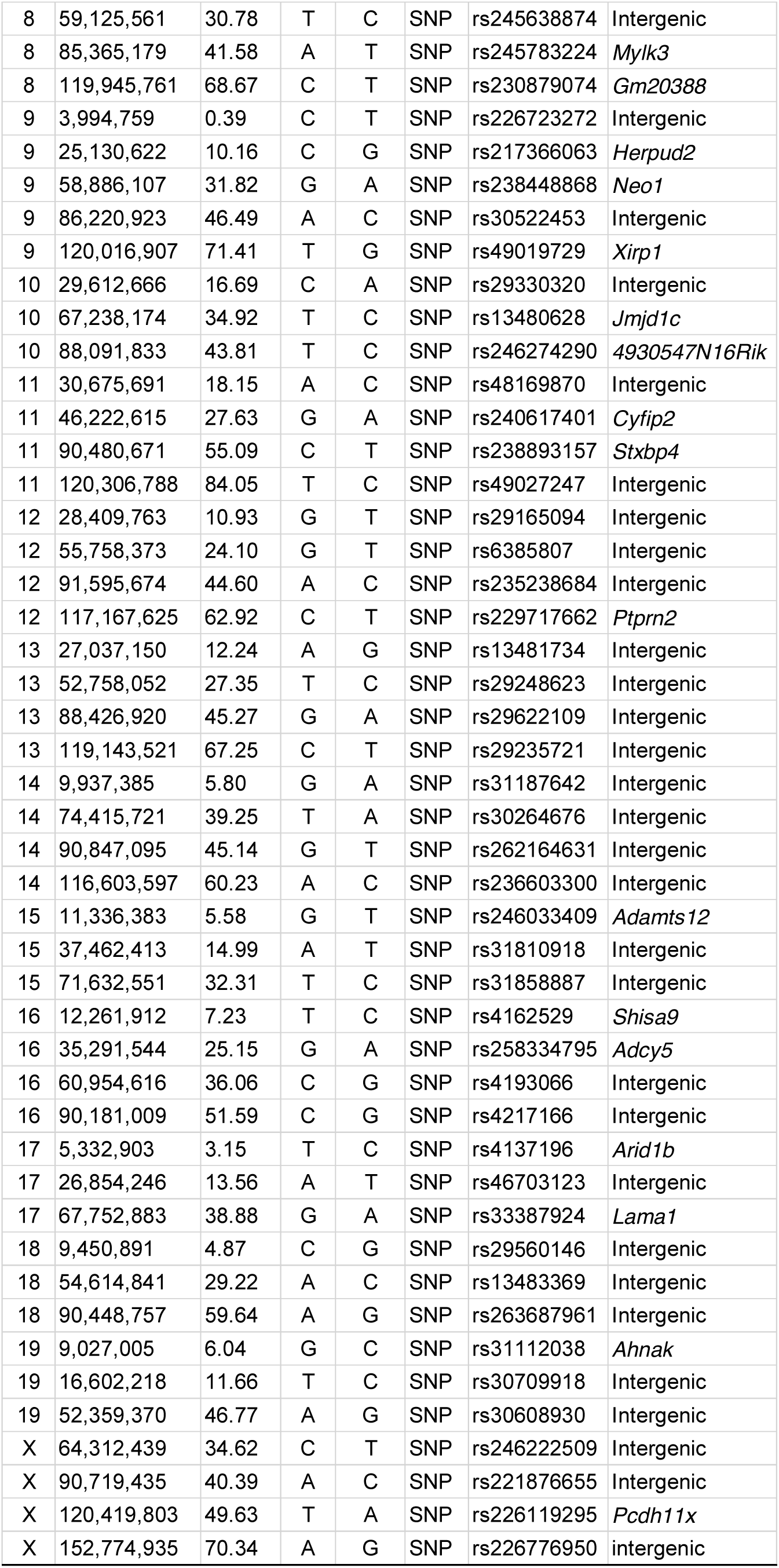
SNP markers for QTL mapping.

**Supplementary Table 2.**
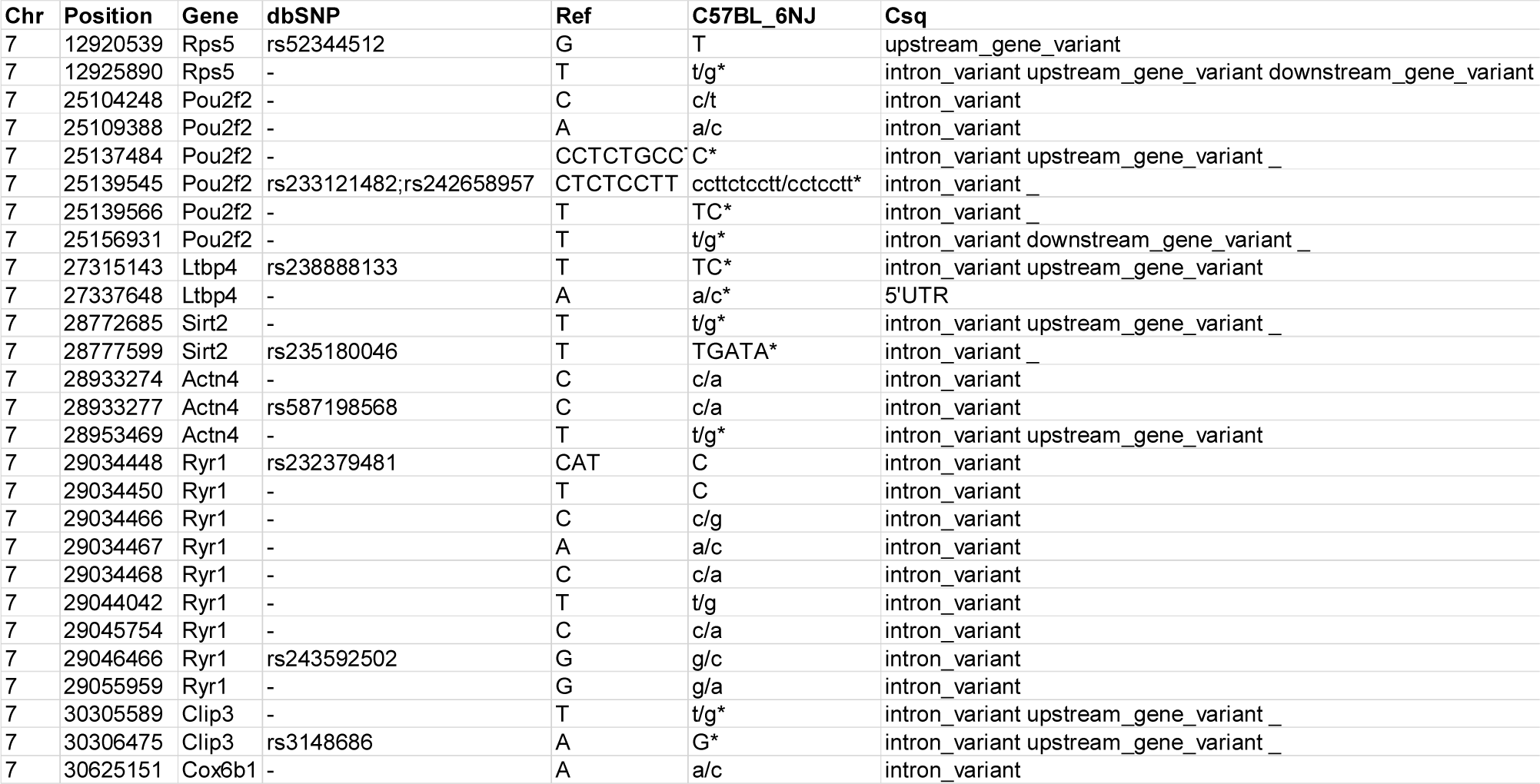
Genes possessing eQTLs within the chromosome 7 hot plate QTL that also possess genetic variants.

**Supplementary Figure 1:**
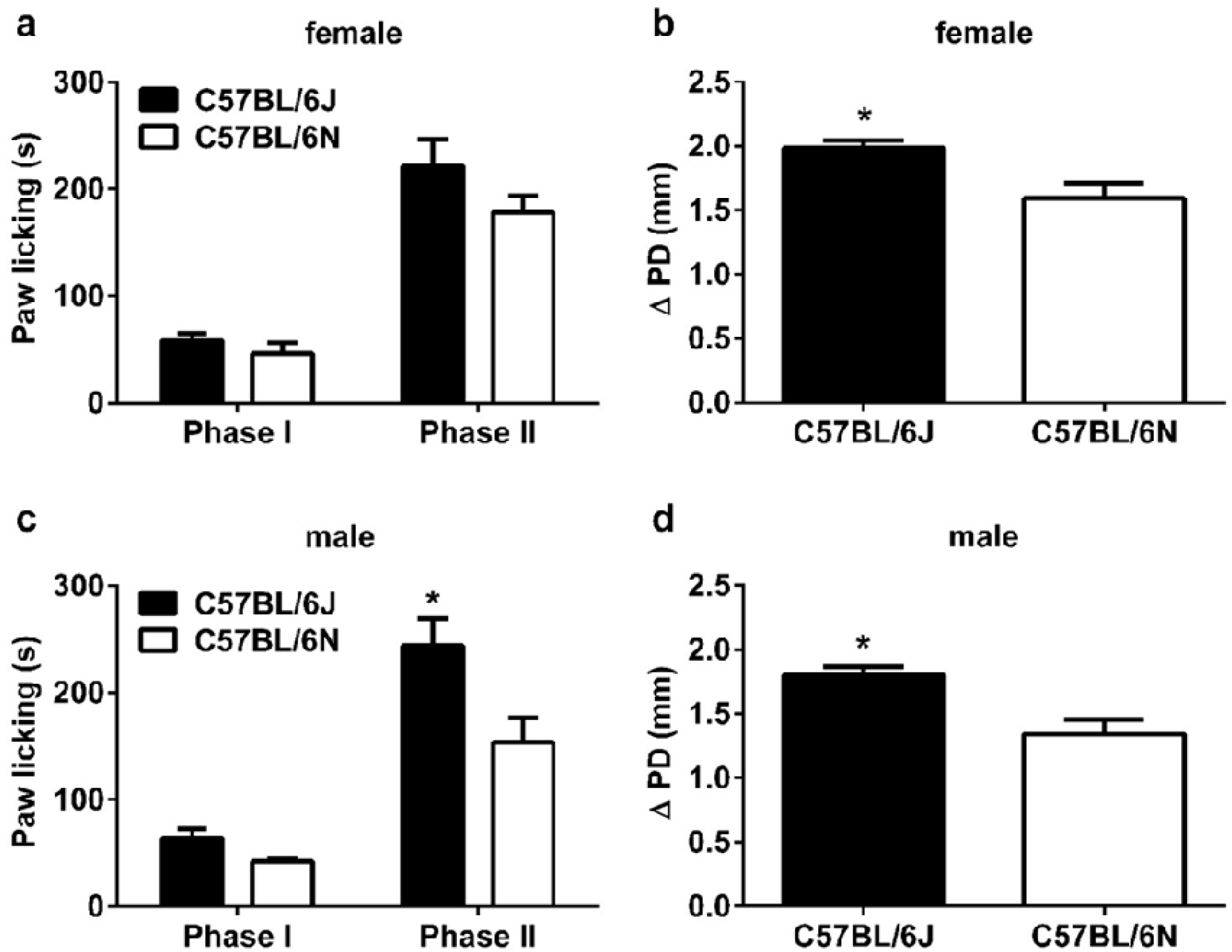
Formalin-induced paw licking behavior and paw edema in female and male C57BL/6J and C57BL/6N mice. The paw licking response after injection of 2.5% formalin concentration into the right paw of C57BL/6J versus C57BL/6N mice. **(A)** female and **(C)** male mice. Changes in paw edema, as measured by the difference in the ipsilateral paw diameter between before versus after injection (ΔPD), in C57BL/6J and C57BL/6N **(B)** female and **(D)** male mice 1 h after injection of formalin. Data are expressed as the mean ± S.E.M. of 6 animals per group. *p<0.05 significantly different from B6J mice.

**Supplementary Figure 2:**
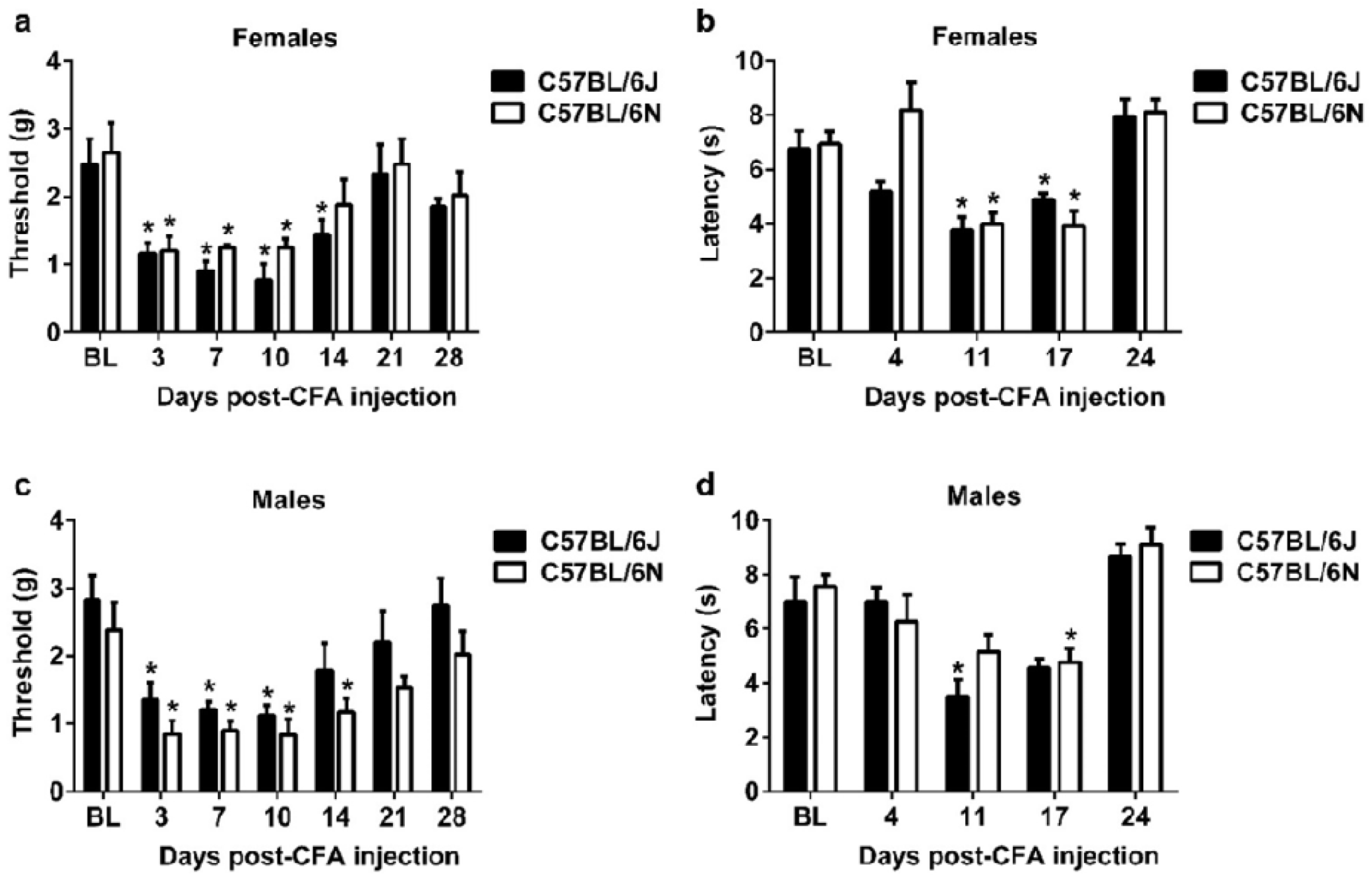
CFA-induced mechanical & thermal hypersensitivity and edema in female and male C57BL/6J & C57BL/6N mice. Differences in mechanical paw withdrawal thresholds in C57BL/6J and C57BL/6N **(A)** female and **(C)** male mice at different days after injection of complete Freund’s adjuvant (CFA, 10 % solution/20 µl). Differences in paw withdrawal latencies (δ PWL=contralateral–ipsilateral hindpaw latencies) in C57BL/6J and C57BL/6N **(B)** female and **(D)** male mice. Data are expressed as the mean ± S.E.M. of 6 mice per group. *p<0.05 significantly different from baseline (BL).

**Supplementary Figure 3:**
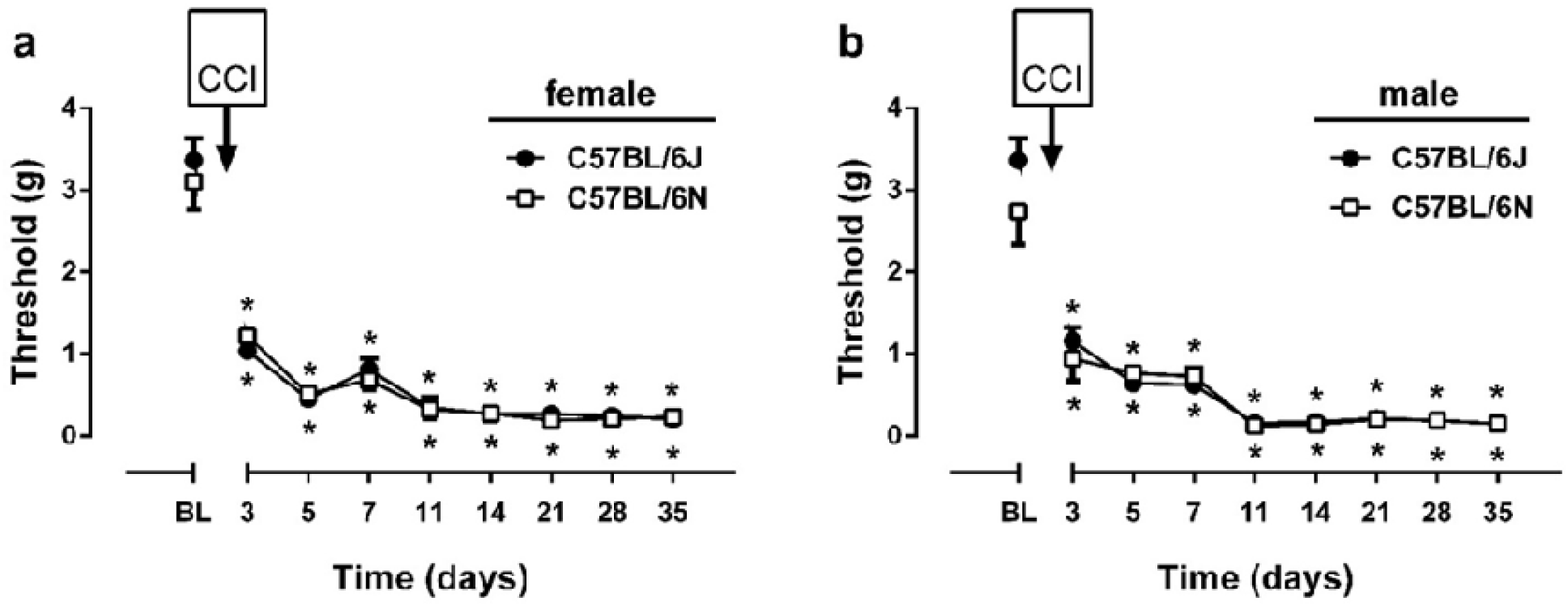
CCI-induced mechanical and thermal hypersensitivity in female and male C57BL/6J and C57BL/6N mice. Differences in mechanical paw withdrawal thresholds in C57BL/6J and C57BL/6N **(A)** female and **(B)** male mice at different times (days) after chronic constrictive nerve injury (CCI) operation. Data are expressed as the mean ± S.E.M. of 6 animals per group. *p<0.05 significantly different from the value of baseline (BL).

**Supplementary Figure 4:**
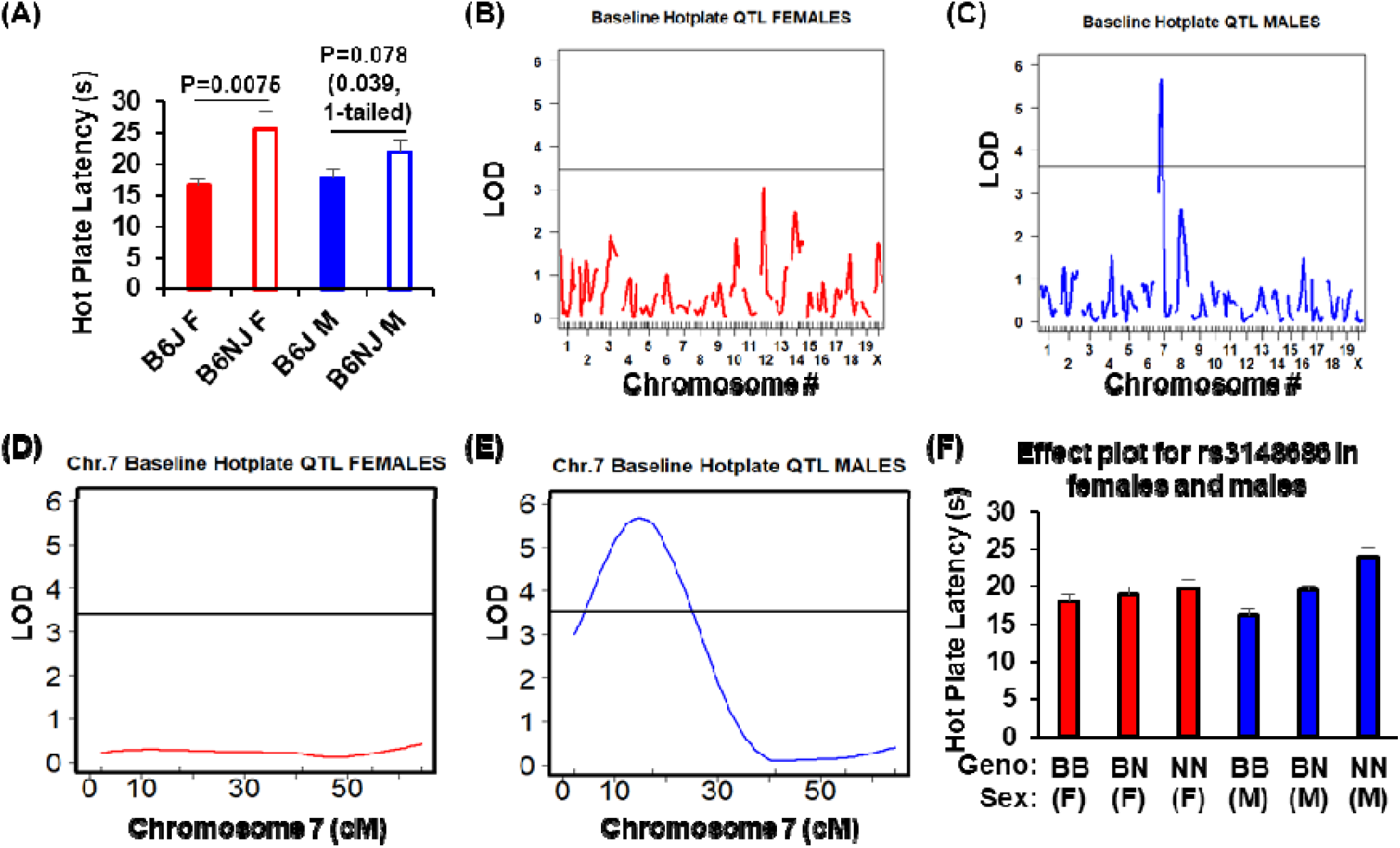
B6 parental substrain differences, QTL plots and effect plots for baseline hot plate latency in female versus male B6J x B6NJ-F2 mice. (A): Female and male B6J mice (n=12 females, 12 male) showed lower hot plate latencies than their B6NJ counterparts (n=11females, 7 males). **(B):** When considering female F2 mice alone (n = 78), no genome-wide significant QTL was detected for hot plate sensitivity **(C):** When considering F2 males alone (n = 86), a genome-wide significant QTL was detected on chromosome 7. **(D):** Chromosome 7 LOD plot is shown for F2 females. **(E):** Chromosome 7 LOD plot is shown for F2 males. LOD = 5.67; peak = 15.30 cM (26.64 Mb); Bayes: 8.30-20.30 cM (14.78 Mb – 34.61 Mb); **(F):** Effect plot at peak associated marker (rs3148686; 31 Mb) for females (F, red bars) and males (M, blue bars). Geno = genotype at rs3148686. BB = homozygous for B6J; BN = heterozygous; NN = homozygous for B6NJ. Data for panels A and E are expressed as the mean ± S.E.M. Horizontal lines for panels B through E indicate the significance threshold (1000x permutations).

**Supplementary Figure 5:**
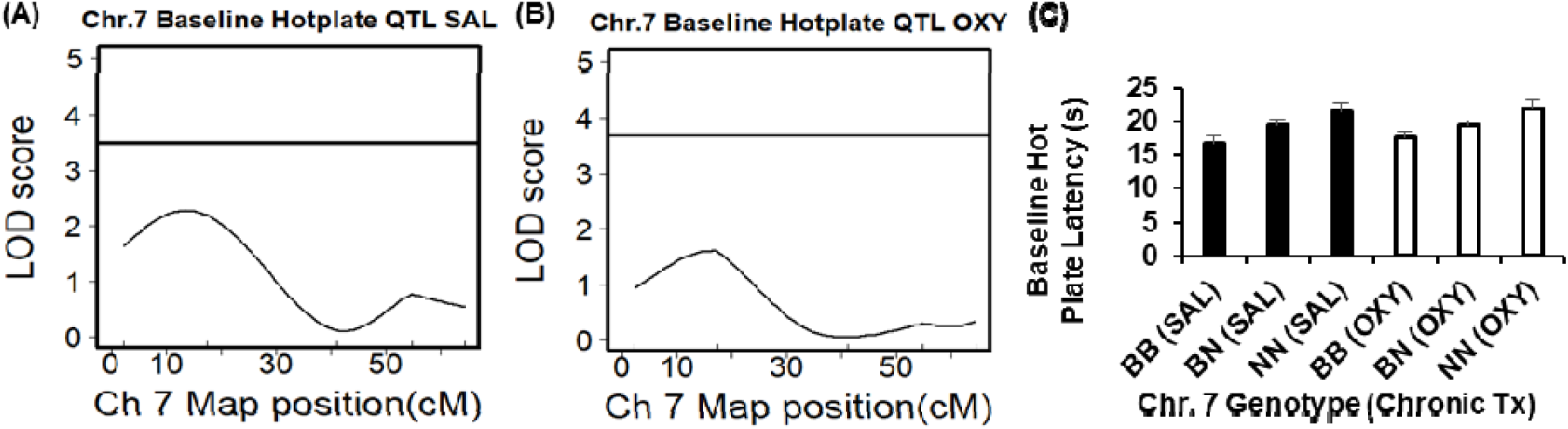
QTLs and effect plots for hot plate latency in B6J x B6NJ-F2 mice with a prior history of repeated SAL versus OXY injections. **(A,B):** Qualitatively similar, trending but non-significant QTL plots were observed, regardless of whether mice had a prior history of SAL injections (**A;** n = 83) or OXY injections (**B**; n = 81; see Methods for details on treatment history). **(C):** Corresponding to the similarly trending QTL plots, qualitatively similar decreases in hot plate latency were observed in mice with a prior history of SAL injections (black bars) versus mice with a prior history of OXY injections (white bars) injections. Mice with two copies of the B6J allele (BB) showed a lower hot plate latency (s) than mice with two copies of the B6NJ allele (NN). Heterozygous mice (BN) were intermediate. Data are expressed as the mean ± S.E.M.

